# FFPE DNA shows two major error profiles derived from deamination of cytosine and methylcytosine that can be mitigated using distinct repair strategies

**DOI:** 10.1101/2023.03.02.530819

**Authors:** Lixin Chen, Minyong Chen, Dan Heiter, Jim Samuelson, Tom Evans, Laurence Ettwiller

## Abstract

Avoiding damage-induced sequencing errors is a critical step for the accurate identification of medium to rare frequency mutations in DNA samples. In the case of FFPE samples, deamination of cytosine moieties represents a major damage resulting in the loss of DNA material and sequencing errors. In this study, we demonstrated that, while damage from deamination of both cytosine and methylated cytosine moieties results in elevated C to T transition, the error profiles and mediation strategies are different and easily distinguishable. While damage-induced sequencing errors from cytosine deamination is driven by the end-repair step commonly used in NGS workflow, DNA damage resulting from deamination of methylated cytosine is another major contributor to sequencing errors at CpG sites. Uracil DNA glycosylase and human thymine DNA glycosylase can respectively eliminate and mitigate both damages in FFPE DNA samples, therefore increasing sequencing accuracy notably for the identification of moderate allelic frequency variants.

## Introduction

Damage has been recognized to be a major limitation for sequencing and accuracy of ancient DNA (1) and recently, several studies have identified artifactual damage in contemporary DNA samples as a pervasive source of sequencing errors. The spectrum of damage on the DNA is heavily determined by the type of handling and storage of DNA samples. For example, acoustic shearing during library preparation introduces oxidative damage of the guanine leading to an erroneous G to T transversions (2) (3). In the oncology field, tissue or biopsy samples are often fixed in formalin-based solutions before embedding in paraffin wax resulting in formalin-fixed, paraffin-embedded (FFPE) tissue blocks. Because FFPE preserves the morphology and cellular details of tissue samples, it is the gold standard method for preservation of human tissues. Unfortunately DNA extracted from FFPE samples is often fragmented, deaminated and contains an elevated amount of abasic sites (4). Despites those damages, studies advocate for optimization of sequencing protocols of FFPE samples rather than seeking to use different biospecimen formats (5).

Artifactual damage to the DNA can be classified depending on the behavior of the DNA polymerase used in sequencing or amplification of the library to either stall at the damage site (blocking damage) or incorporate the wrong base (mutagenic damage). This distinction is important because only mutagenic damages are responsible for sequencing error and the usage of different families of polymerases during library preparation can result in the conversion of a blocking damage to mutagenic damage and vise-versa. The usage of a dU-bypass DNA polymerase leads to the conversion of a dU from blocking to mutagenic leading to C to T transition, an attribute largely used for bisulfite sequencing (6).

Thus, most of the polymerases used in sequencing contain a dU pocket that prevents amplification of dU containing fragments (7). A notable exception is the end-repair, a crucial step in the ligation-based library preparation that consists of filling-in overhanging 5’ end uses T4 DNA polymerase, an enzyme of a different family of polymerase known to bypass dU damages (8).

In this work, we analyzed the effect of end-repair on the rate of sequencing errors for FFPE samples and demonstrated that end repair can permanently fix damage by incorporating the mispaired nucleotide on the opposite strand of damage. We also found that the characteristic of the polymerase in the end-repair step allows the conversion of a blocking damage to a mutagenic damage leading to further sequencing errors with additional mutagenic signatures. We also show DNA damage caused by methylated cytosine deamination in the CpG context of FFPE DNA is the major contributor to damage-induced sequencing errors in FFPE DNA samples. Pre-treatment of FFPE DNA before library construction using Uracil DNA glycosylase and human thymine DNA glycosylase removes damages generated by both cytosine deamination and methylated cytosine deamination, therefore increasing sequencing accuracy.

## Result

### FFPE DNA standard NGS sequencing shows two major CG:TA error profiles

Previous studies have reported that deamination of cytosine to deoxyuracil (dU) is the main damage induced by spontaneous hydrolysis in FFPE treated samples [(9, 10)]. dU is known to pair with adenosine leading to erroneous CG:TA transitions during sequencing (9, 10). Inconsistent with this statement, polymerases used for NGS library amplification and sequencing contain a uracil-binding pocket that effectively stalls the polymerase at dU sites. Thus, dU being a blocking damage, dU containing fragments should not be amplified and sequenced. We examined this apparent contradiction by investigating the cause of the elevated CG:TA transition reported in FFPE samples.

For this, we amplified a library of FFPE-derived genomic DNA using either a polymerase that stalls at dU (Q5 DNA polymerase or Q5) or dU-bypass DNA polymerases (either Q5U® or Taq DNA polymerase). To demonstrate the presence of damage, we also perform these experiments with and without treatment with either DNA damage repair mix (NEBNext FFPE DNA Repair v2 Module) or Uracil-DNA Glycosylase (UDG). DNA damage repair mix is a cocktail of enzymes that repairs various types of DNA damages including those blocking damages such as abasic sites, nicks, gaps, and thymine-dimers and those mutagenic damages such as cytosine deamination and oxidative damages. UDG is included in the DNA damage repair mix and catalyzes the hydrolysis of the N-glycosidic bond from deoxyuridine, thus is specific to dU damage (11).

To observe the effect of damage, we calculated the number of CG:TA transitions with high quality scores (Q-score > 30 to eliminate most of the sequencing errors) and normalized this number to the sequencing depth at cytosines for comparison (**Material and Methods**). As expected, we found that the CG:TA transition is the highest for genomic DNA amplified using dU-bypass DNA polymerases (**Figure 1A**). Importantly, we also found an elevated CG:TA transition in the Q5 Amplified samples (**Figure 1A**). DNA repair treatment using FFPE DNA Repair v2 Module further lowers the CG:TA transition in all samples suggesting that the excess of CG:TA in non-DNA repair treated samples is due to damage of cytosines leading to dU. UDG treatment lowers the CG:TA transition in Q5 amplified samples to levels comparable to repaired samples (**Figure 1A**) demonstrating that the excess of CG:TA transition in the Q5 amplified samples is due to the presence of dU. These results are consistent with the idea that FFPE treatment leads to erroneous CG:TA transitions but inconsistent with the usage of a polymerase containing a uracil-pocket that stalls at dU containing DNA.

**Figure 1.**
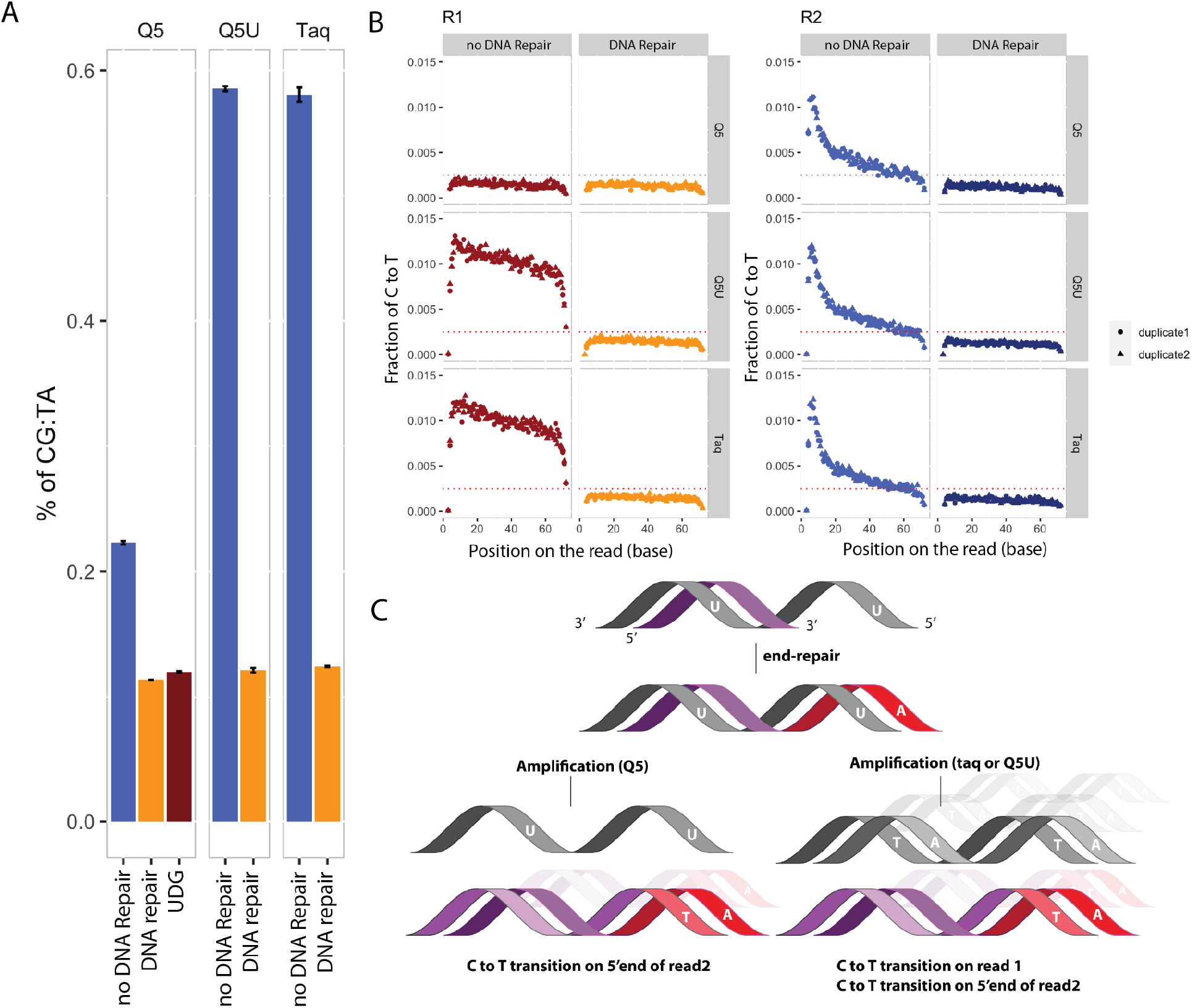
Effect of polymerases on the accuracy of sequences in FFPE samples. **A**. Overall percentage of CG:TA transitions in sequencing reads after library amplification using Q5 (left), Q5U® (center) and Taq DNA polymerase (right), with (DNA repair (orange), UDG (brown)) and without (blue) DNA repair treatment of the lung tumor FFPE sample. Experiments were done in duplicate and error bars are reflecting the variability between replicates. **B**. Profiles of C to T transitions on read 1 (R1, left) and read 2 (R2, right) as a function of read position (in bp). Libraries were amplified using Q5 (top), Q5U® (middle) and Taq DNA polymerase (bottom). Experiments were done in duplicate. Fraction of C to T is a global count of the of C (on the reference genome) to T (on the mapped reads) normalized to the total number of C to C matches. **C**. Schema representing the fate of the deaminated strand (gray) and the un-damaged strand (purple) after end repair (red) and amplification with Q5 (left) and Taq or Q5U (right).

To further investigate the origin of the elevated CG:TA notably for Q5 amplified samples, we took advantage of the fact that paired-end sequencing using Illumina Y shaped adaptors ‘orients’ the direct readout of mutagenic damages on the first of a paired-end read corresponding to read 1 (R1) (10). We therefore subjected the sequencing result to our orientation-aware variant calling algorithm and investigated the profile of C:T and G:A transitions separately at various read positions. An overall elevation of C:T transition is observed on R1 specifically for libraries amplified using Q5U® and Taq DNA polymerase (**Figure 1B**,**C**) consistent with the polymerase dU bypass properties. Conversely, no apparent elevation of C:T transition is observed in R1 for the Q5 libraries consistent with polymerases with dU pockets. In fact, the level of C to T transition in R1 for Q5 libraries is comparable to the one observed after DNA repair (**Figure 1B**,**C**). Instead, the elevation of C to T transition in these samples is observed for read 2 (R2) with a stark bias towards the 5’ end of the read accounting for the CG:AT elevation observed in **Figure 1A**. Furthermore, this 5’ end C to T excess is independent of the polymerase used for amplification and is eliminated when the DNA is repaired prior to library construction. From the C to T error profile bias towards the 5’ end, we hypothesize that this elevation of C to T transition on read 2 is due to the end-repair, a necessary step during ligation-based library construction (**Supplemental figure 1A**). Indeed, this step involves a polymerase with dU bypass properties, T4 DNA polymerase filling 3’ single stranded DNA. Because T4 DNA polymerase does not stall at dU (8) an A opposite of a dU base is incorporated on the complementary strand. The complementary strand is amplified and sequenced leading to an G to A transition (or a C to T transition on read 2). Additionally, a previous study has demonstrated that between 32–57% of bases could be resynthesized *in-vitro* during the end repair step for FFPE samples (12), indicating that the filling during end-repair can provide a notable source of sequencing errors.

**Supplemental Figure 1:**
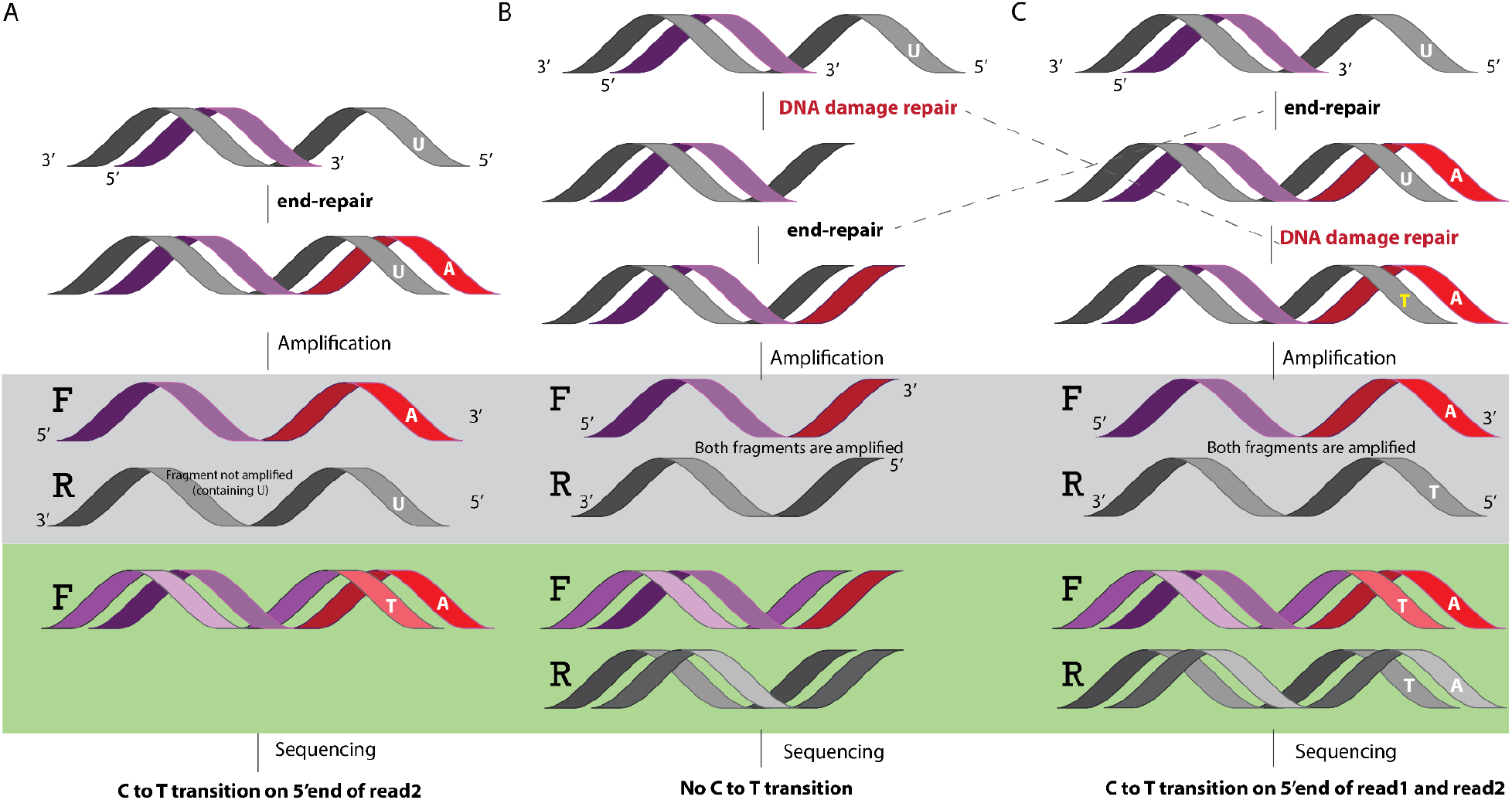
End-repair, DNA damage repair and sequencing accuracy. **A**. How the end-repair step turns deamination damages into artifactual C to T error : Schema describing the contribution of the end-repair step in fixing a dU damage in single stranded DNA (5’ overhangs) towards a C to T sequencing error. The end-repair strand (F, forward) is sequenced while the reverse strand (R) cannot be amplified and sequenced because it contains a U. **B**. Effect of DNA damage repair performed before end-repair : DNA damage repair removes dU on single stranded DNA before end repair. **C**. Effect of DNA damage repair performed after end-repair : DNA damage repair fixes the error introduced during end-repair and allows the sequencing of both the fragments that originally contained a U (R, reverse) and the end-repair strand (F, forward). Thus, performing DNA damage repair after end-repair leads to more C to T errors.

### The end-repair step is one major driver for damaged-induced sequencing error

Several lines of evidence confirm the hypothesis that the elevation of C to T transitions observed on read 2 is resulting from the end-repair step. First, the use of Uracil-DNA Glycosylase (UDG) prior to library preparation and sequencing shows that the elevated C to T error observed on read 2 vanishes after UDG treatment (***Figure 1A***). Similar results are obtained if DNA damage repair mix is used instead of UDG (***Figure 1A***). These results demonstrate that the elevated C to T transition is the result of cytosine deamination. Next, the amplification of FFPE samples with a polymerase containing a mutated dU pocket shown to bypass dU identifies an excess of C to T uniformly across read 1 (***Figure 1B***). This result clearly demonstrates the presence of a large degree of damage that leads to C to T erroneous variants in R1 sequence. Given the property of the polymerase to bypass the dU damage, the responsible damage is very likely deamination of cytosine. Interestingly, while it was speculated that the rate of deamination of cytosine is elevated in single stranded DNA (13), we observed an almost uniform distribution of C to T variant rate in the R1 sequences suggesting that the deamination is more or less equally distributed in single and double strand DNA.

Finally, the most direct evidence of the effect of end repair on fixing the damage comes from performing a comparative analysis using DNA damage repair mix before and after end repair (**Supplementary Figure 1B and C)**. We show that the CG:AT error rate elevation is observed for both unrepaired samples and samples for which DNA damage repair is performed after end-repair. Counterintuitively, the CG:AT elevation is the highest if DNA damage repair is performed after end-repair (**Figure 2A**). This result can be explained by the fact that the enzymes in the DNA damage repair mix will remove uracil at filled sites but replace the damaged base with a thymine instead of a cytosine. This is because the template DNA is an adenosine wrongly added during the end repair step (see **Supplementary Figure 1** for explanation). Consistent with this scenario, we observe an elevated C to T variant in read 2 being mirrored by an elevated C to T variant frequency in read 1 when DNA damage repair mix is applied after end repair (**Figure 2B)**.

**Figure 2.**
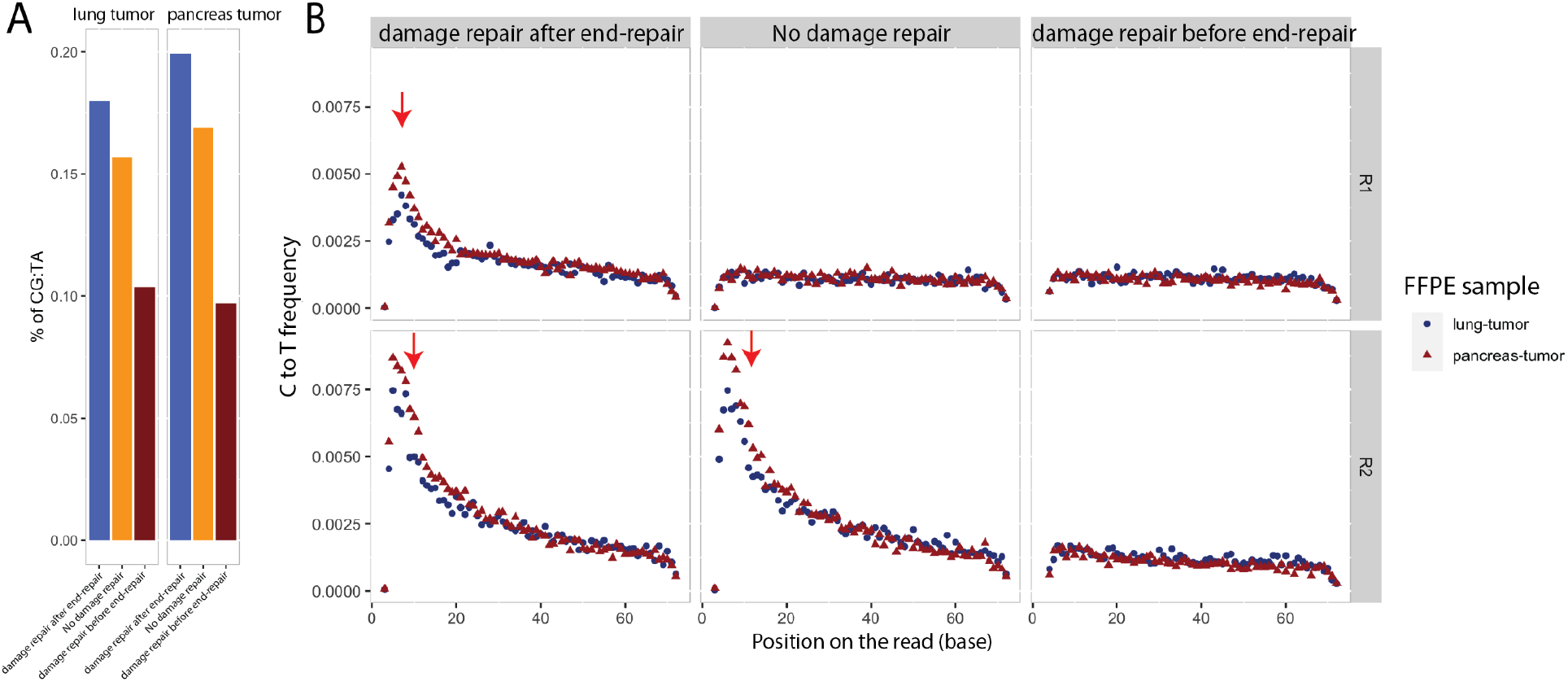
Contribution of end-repair on CG:TA transition. **A**. Overall percentage of CG:TA transitions in sequencing reads for lung (left) and pancreatic (right) tumor FFPE samples. Samples were either left untreated (orange) or treated with DNA repair mix before (red) and after (blue) end repair. **B**. Breakdown of the fraction of C:T transitions according to Read types, Read 1 (R1, top) and Read 2 (R2, bottom) and position on the read starting from the 5’ end. Two FFPE samples have been used for these experiments : Lung tumor FFPE sample (Blue) and pancreatic tumor FFPE sample (red) treated with DNA repair after end repair (left), before end repair (right) and no treatment (middle). Fraction of C to T is a global count of the of C (on the reference genome) to T (on the mapped reads) normalized to the total number of C to C matches. Red arrows indicate positions of increased C to T transition rate.

Collectively, these results show that a notable fraction of the artifactual CG:TA transitions previously observed in FFPE treated samples are, in fact the result of a mis-incorporation of a dA opposite to the dU containing strand by the T4 DNA polymerase during the end-repair step of library preparation.

### Deamination of 5mC is another driver for damaged-induced sequencing error in FFPE samples

Elevation of the C to T error rate in FFPE samples cannot completely be explained by mis-incorporation of dA at dU damages during the end-repair step. Deamination of methylated cytosines (5mC) leads to a thymine base (dT) that, contrary to dU, is a regular base for which all DNA polymerases will incorporate a dA instead resulting in a C to T error. Thus, the profile of C:T transition at 5mC sites is expected to be distinct from the C:T transition observed at non methylated sites. Since most of the methylation happens at CpG sites in human, we performed Illumina sequencing of an FFPE sample (lung normal tissue, Biochain) after Clear seq comprehensive cancer panel enrichment (Agilent Technologies) and examined the C:T transition in CpA, CpT, CpC and CpG context separately. While the C:T transitions in CpA, CpT, CpC follows the expected profile consistent with the end-repair step incorporating the wrong base (see previous section), elevation of the C:T transition in read 1 is only observed for the CpG context and is uniformly distributed across the reads (**Figure 3a**). Furthermore, DNA repair using FFPE DNA Repair v2 Module (that cannot repair a G:T mispairing) does not lower down the rate of C:T transition in read 1 (**Figure 3a**). This data suggests significant deamination of 5mC to T in FFPE samples leading to C:T transition.

**Figure 3.**
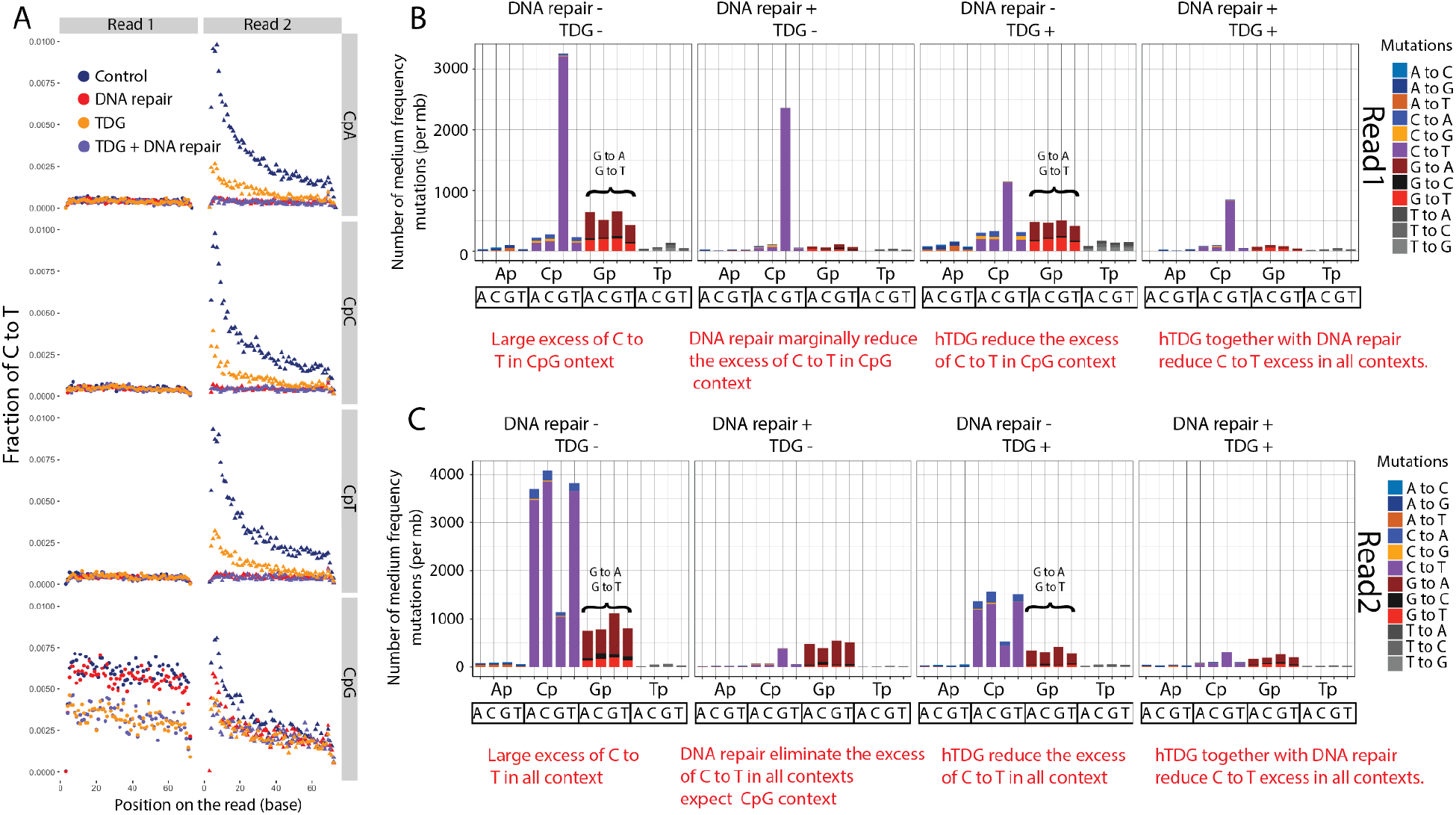
**A.** Profile of C:T transition according to Read types, Read 1 (left) and Read 2 (right) and position on the read for CpA (top), CpC (mid-top), CpT (mid-bottom) and CpG (bottom) contexts. The sample is an FFPE treated normal lung hDNA paired-end sequenced after Clear seq comprehensive cancer panel enrichment (Agilent). Exome sequencing : B and C. Profiles of medium (1-10%) allelic frequency errors in various contexts for read 1 (**B**) and read 2 (**C**). Sequencing depth has been downsample to 300-fold coverage for comparison, thus the variant read number for 1-10% allelic frequency is between 3 and 30 reads supporting the presence of a variant (with Phred Quality Scores Q>30). The excess of G to T can be attributed to oxidative damage.

### hTDG and MBD4 are effective enzymes in removing deaminated 5mC and 5hmC

To demonstrate the hypothesis of significant deamination of 5mC to T in FFPE samples, we evaluated two DNA Glycosylases; human thymine DNA glycosylase (hTDG) and mouse methyl-CpG binding domain protein 4 (MBD4), more specifically, hTDG and the C-terminal mismatch-specific thymine glycosylase domain of MBD4 (15). For hTDG, we either used hTDG WT or hTDG-A145G, a TDG variant that exhibits 13-fold increase activity in G:T mispairing repair compared with native TDG (14). hTDG and MBD4 have previously been shown to remove thymine moieties from G/T mismatched pairs by hydrolyzing the carbon-nitrogen bond between the sugar-phosphate backbone of DNA and the mispaired thymine.

Fragments containing these abasic sites will not be amplified and thus, the C to T error rate from deamination of 5mC is expected to decrease.

We first characterized human thymine-DNA glycosylase hTDG repair on G:T and G:hmU mismatch activity on various sequence contexts using a NGS-based assay. For this, we used fully modified XP12 and T4gt phage genomic DNAs that harbor 5mC (16) and 5hmC (17) respectively and were subjected to a limited deamination using heat alkaline treatment following similar conditions as previously published (18) with some minor changes as described in **Material and Methods**. Using such treatment, the deamination rate of XP12 has been previously shown to be even across all contexts (18). We further demonstrate that the deamination rate of 5hmC is proportional to the heat alkaline treatment time (**Supplementary Figure S2A**) and evenly distributed across all analyzed contexts (**Supplementary Figure S2B**).

Deamination of 5mC is leading to G/T mispairing while 5hmC is leading to G/hmU mispairing. A decrease in the C to T error rate in the TDG treated samples compared to no TDG treatment is indicative of the enzyme activity. We showed that hTDG treated samples compared to untreated samples have a lower C to T error rate in read 1 for both damaged XP12 and T4gt genomic DNA indicating glycosylase activity for both G/T and G/hmU mispairing is consistent with previous studies using oligo-based assay (19).

Nonetheless, the C to T error rate is above the baseline obtained for undamaged XP12 and T4gt indicative of partial activity only, likely the result of context specificity. Consistent with this hypothesis, the decrease of C to T error rate is only observed in CpG and CpA context for Xp12 indicating that (**Figure 4**) the removal of the thymine moieties from G/T mispairing is specific to CpA and CpG contexts. Conversely, for T4gt, the decrease of C to T error rate is observed in all contexts indicating that the removal of the thymine moieties from G/hmU mispairing is unspecific. Quantification shows that TDG for G/T mispairing has CpG>CpA preferences as previously shown (20). The context preference for G/hmU mispairing is CpG>CpA>CpC>CpT.

**Figure 4.**
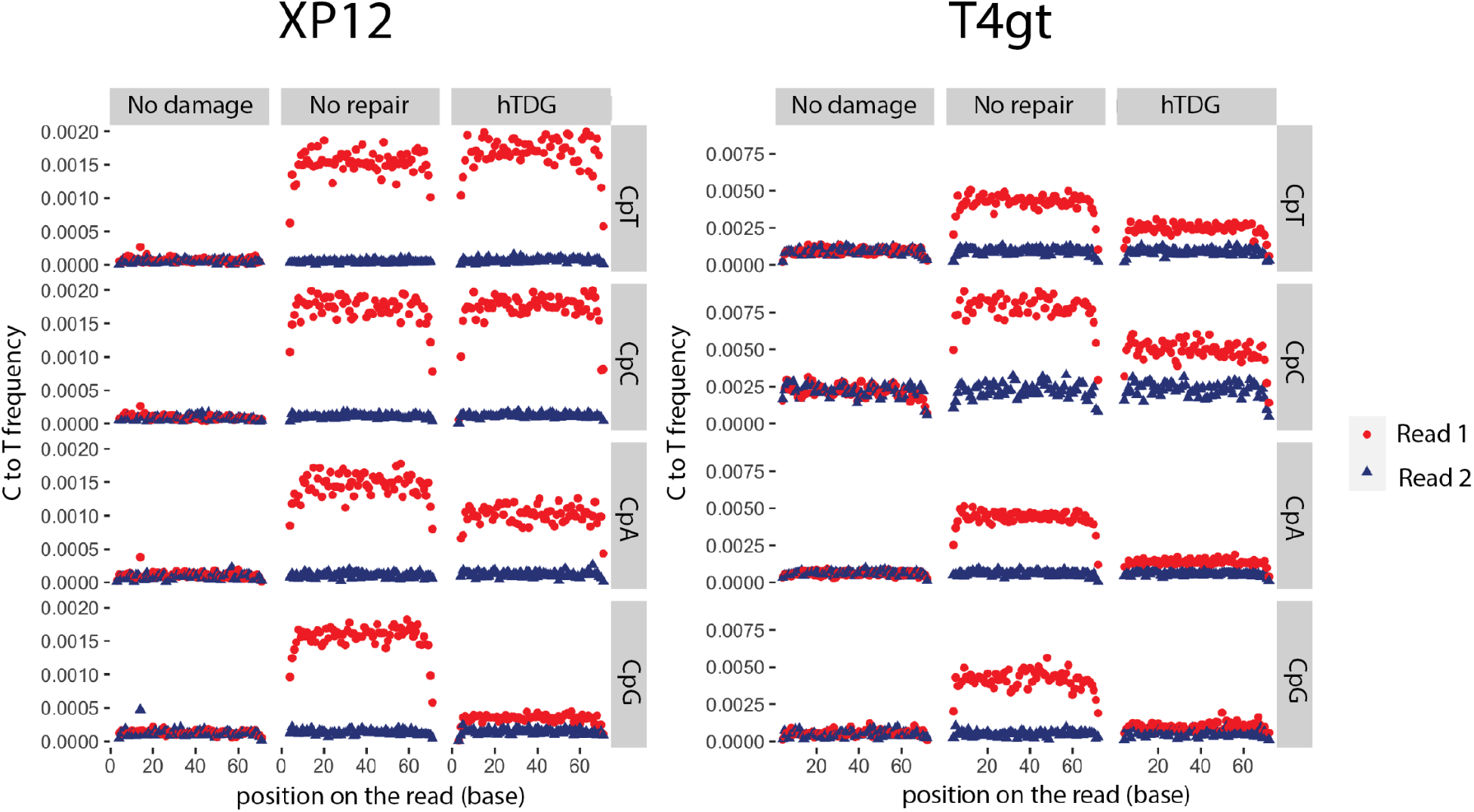
Frequency of C to T error for XP12 genomic DNA containing 5mC (left) and T4gt genomic DNA containing 5hmC (right) for read 1 (red) and read 2 (blue) at CpT (top), CpC (center-top), CpA (center-bottom) and CpG (bottom). Samples (No repair and hTDG) have been damaged using heat alkaline treatment (20min) and subsequently repaired with hTDG (data shown for hTDG-A145G). A control with no damage (No damage) has been included in the experiment to evaluate the baseline sequencing error.

We also perform similar experiments, replacing hTDG with MBD4 (15). MBD4 belongs to the helix-hairpin-helix (HhH) DNA glycosylase superfamily and has been shown to catalyze the removal of T, U and hmU paired with guanine (G) (21) (22). MBD4 decreases the C to T error rate on both XP12 and T4gt indicating that the enzyme is active on both G:T and G:5hmU (**Supplementary figure 2B**).

Interestingly, the activity can be observed in all sequence contexts for both hmC and 5mC deamination damages (**Supplementary Figure 2B**). Because hTDG showed stronger activity in the CpG context, we selected this enzyme for the subsequent experiments on human genomic DNA.

**Supplemental Figure 2:**
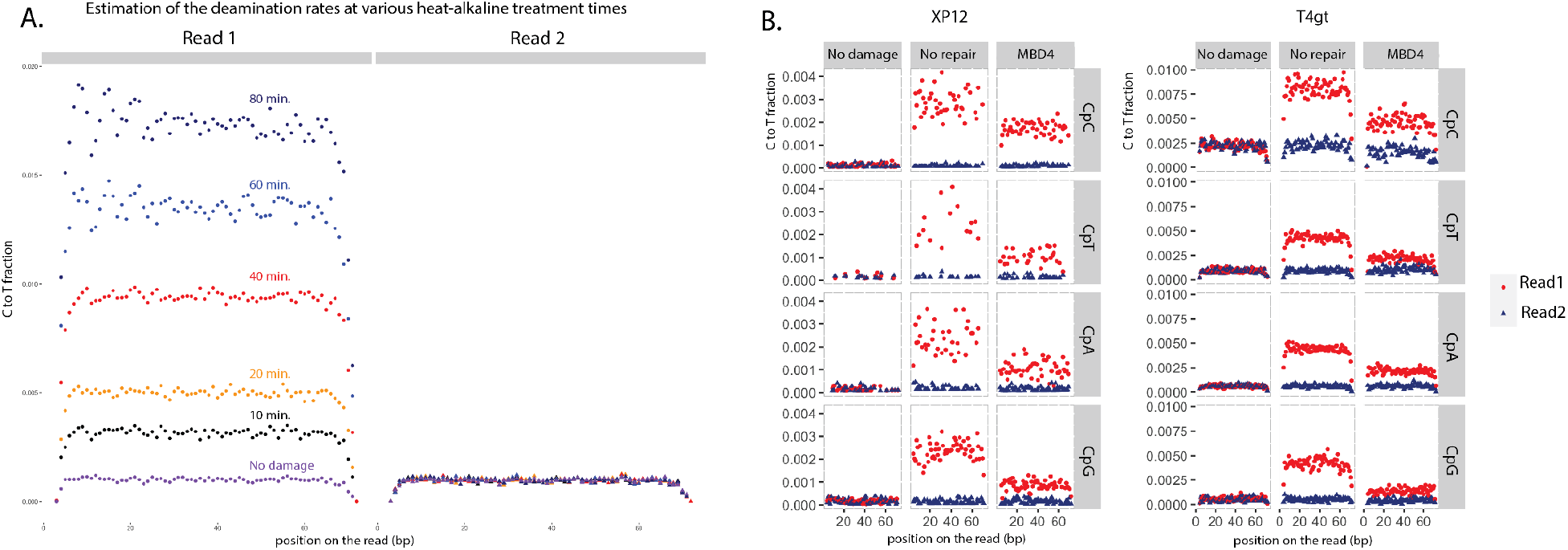
Deamination of modified cytosine moieties. **A**. Deamination time-course of T4gt genomic DNA (5hmC). Heat alkaline treatment of the genomic DNA has been performed for 0 (no damage, purple), 10 (black), 20 (orange), 40 (red), 60 (blue) and 80 minutes (dark blue). The fraction of C to T transversions were plotted according to position on the read (starting from 5’ end) and read 1 (left) and read 2 (right) of the paired-end reads. **B**. Fraction of C to T transversions in all 4 contexts (CpC, CpT, CpA and CpG) of XP12 (5mC, left) and T4gt (5hmC, right) that has been deaminated using heat alkaline treatment for 20 minutes or not (No damage, left panel). Deaminated samples were either not repaired (middle panel) or repaired with MBD4 (right panel).

### Deamination of 5mC is also the cause of C to T errors in FFPE samples

We treated the FFPE sample with hTDG and found that this treatment lowered the rate of C:T transition in read 1 in CpG context (**Figure 3a)**. Conversely, treatment with DNA damage repair mix did not change the rate of C:T transition in read 1 compared to the untreated sample (**Figure 3a)**. These results indicate that the elevated C to T rate in read 1 is due to deamination of 5mC.

### Deamination of both cytosines and methyl cytosines account for the majority of the medium-frequency mutation in FFPE samples

Most of the applications involving sequencing of FFPE samples do not use the raw C to T transitions on reads but rather focuses the identification of either high allelic frequency variants for which a large fraction of reads show evidence of non-reference nucleotide (such as germline mutations) and/or medium to low allelic frequency variants (such as somatic mutations) for which a minority of reads show evidence of a non-reference nucleotide at any particular genomic position. We therefore investigated the effect of C and 5mC deamination in the identification of low (<1%), medium (1-10%) and high (> 10%) allelic frequency substitution. To achieve the sequencing depth necessary for such an analysis, we performed a targeted genome sequencing using the Clear seq comprehensive cancer panel enrichment (Agilent) on an FFPE sample (Lung normal) that has been treated or not with DNA repair and/or hTDG (WT). To obtain defined variant read numbers across samples, the sequencing depth has also been set to 300-fold coverage for both read 1 and read 2 respectively.

As expected, a total of 143,171; 23,383 and 1,044 positions per mb show low, medium and high allelic frequency substitution in the control untreated FFPE sample respectively. In contrast, the same material treated with DNA repair and hTDG shows 93,454; 3,226 and 1,122 positions per mb with low, medium and high allelic frequency substitution respectively (**Supplementary Table 1**). These results indicate that, while the majority of substitutions can be observed at low frequency, the major effect of damage is observed for the medium (1-10%) frequency variants with 7.2 fold increase in the substitution level attributable to deamination damage.

Breaking down the medium frequency variants by substitution spectrum (all transitions and transversions), reads (read 1 and Read 2) and contexts (XpT, XpC, XpA and XpG, with X being the substituted A,T,C or G), we observed [1] a large excess of C to T transition in CpG context on read 1 that can be mitigated by hTDG treatment [2] a large excess of C to T transitions in all contexts on read 2 that can be mostly eliminated using DNA repair and [3] G to A transition in all contexts on read 1 that can be mostly eliminated using DNA repair (**Figure 3B**). These observations are in agreement with the previous result that FFPE-induced deamination damages of both C and 5mC are the leading cause of medium frequency variants in the untreated samples. Similar observations, albeit less substantial, can be made on the low frequency substitution (**Supplementary Figure 3A**). In contrast, high frequency substitution closely reflects genome evolution at the human population level with a large fraction of balanced CG:TA and TA:CG transition. Furthermore, the spectra is not altered by hTDG and/or DNA repair treatment indicative of true germline variations.

**Supplemental Figure 3:**
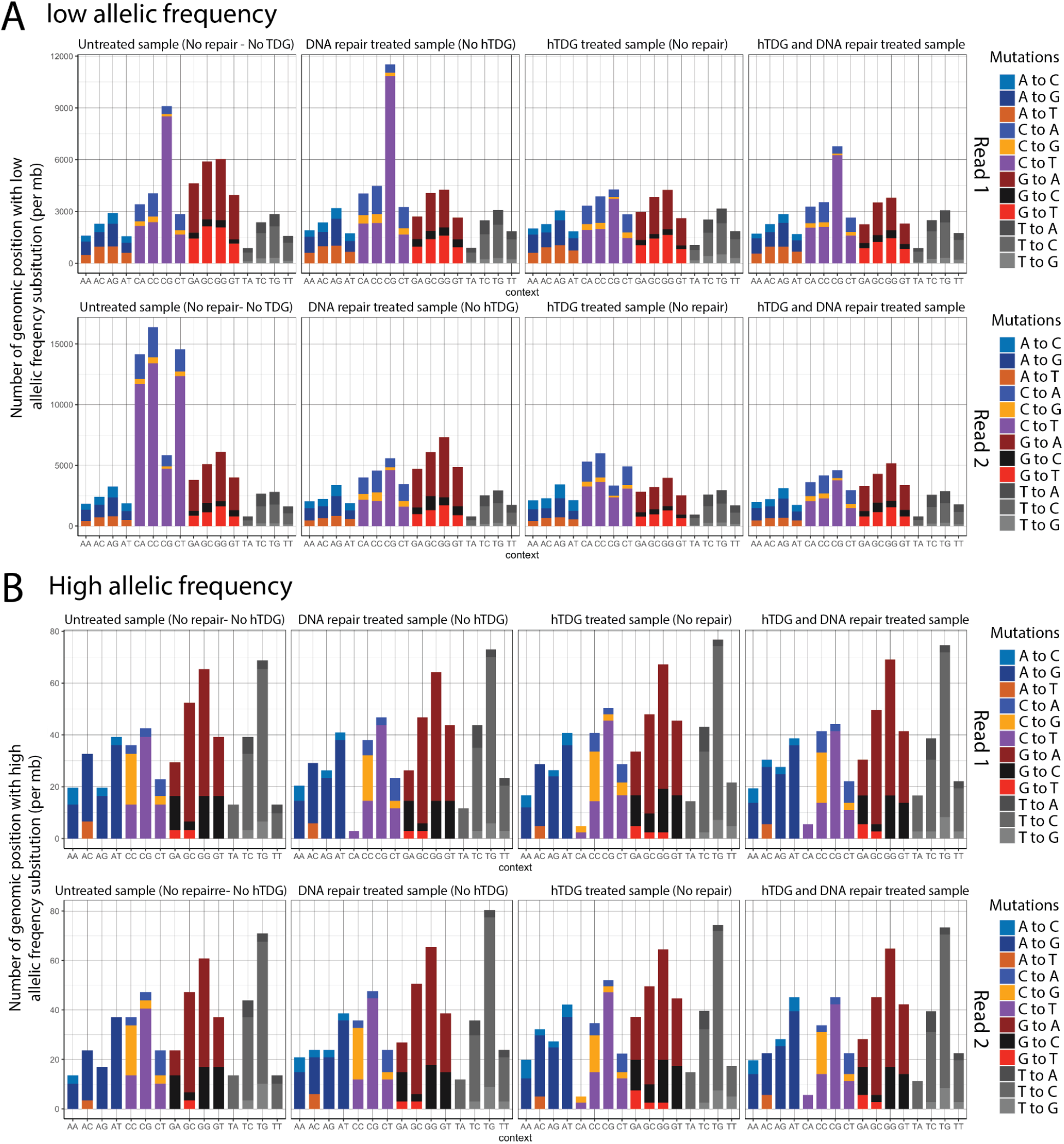
Substitution profiles for low (less than 1% **A**) and high (more than 10% **B**) allelic frequency substitution. Barplot representing the number of genomic positions per megabases (mb). The barplots are color coded according to the substitution types and organized by contexts (XpA, XpT, XpC and XpG with X= A, C, T or G) and reads (read1, top and read2, bottom). Sequencing depth has been downsample to 300-fold coverage for comparison, thus the variant read number for high (>10%) allelic frequency is above 30 reads supporting the presence of a variant and low (<1%) allelic frequency is below 3 reads supporting the presence of a variant. The equivalent result for medium (1 to 10%) allelic frequency substitution is plotted on Figure 3B and C. All reads substitution have a Phred Quality Scores Q>30. CpA context has no high allelic frequency substitution in Read 2 in the DNA repair sample. The excess of G to T can be attributed to oxidative damage.

## Discussion

In this study, we demonstrated that deamination damage of genomic DNA can lead to two very distinct C to T error profiles depending on whether the deamination affects cytosine or 5-methylcytosine moieties. Deamination of cytosines leads to sequencing errors introduced during the end repair step of library construction while deamination of 5mC leads to errors introduced during DNA amplification. Consequently, while both deamination events lead to a C to T transition, the error profiles are different : In the case of cytosine deamination, most of the errors occurs at the 5’ end of the second read of the paired-end reads while 5mC deamination leads to C to T errors evenly distributed across the first read of the paired-end reads.

Most of the sequencing errors that are attributed to damage in FFPE samples result in low allelic frequency ranges that are typically filtered out by mutation callers. Importantly, we also show that deamination damages of both cytosine or 5-methylcytosine can lead to rates of sequencing errors that confound the identification of variants with medium allelic frequency ranges (1-10%), frequencies that are expected for somatic mutations in cancer. Conversely, the level of damage is not sufficient to visibly alter the identification of high frequency variants (>10%), at least for the FFPE samples that we analyzed.

These deamination events require two mediation strategies to avoid sequencing errors : deamination of C can easily be repaired using commercially available repair enzymes with a complete recovery of amplifiable DNA and return of C to T error to baseline. Deamination of 5mC requires specialized enzymes such as hTDG that only recognize the deamination event on a double-stranded context and excise the base. hTDG has been shown to reduce sequencing artifacts notably on the EGFR T790M mutations in FFPE samples (23).

As this work focuses on human FFPE samples we used hTDG enzyme to mitigate the 5mC damage. hTDG shows strong preference for CpG and to a lesser extent CpA contexts which correspond to the contexts for which the majority of methylation occurs in humans. We also show that MBD4 effectively mitigates the C to T error in other contexts as well and thus, may be more adapted to other samples such as plant genomic DNA for which methylation occurs in non-CpG contexts.

While this study focuses on demonstrating the origin of C to T errors, the mitigation strategies used in this study can be used to improve sequencing accuracy for samples that have been subjected to various deamination due to improper storage/ treatment of DNA leading to DNA damage. This study also serves as a framework to study the origin of other damages and provides assays for measuring DNA damage and repair in all sequencing contexts.

## Supporting information

Number of low, medium and high frequency variants for various treatment of FFPE samples

## Acknowledgments

We thank the NGS core sequencing group, Jurate Bitinaite and Rita Vaiskunaite for the expression, purification and initial characterization of the hTDG enzyme. Peter Weigele and Yan-Jiun Lee for providing purified XP12 phage genomic DNA.

## Conflict of interest statement

LC, MC, DH, and JS, TE and LE are employees of New England Biolabs, Inc, a manufacturer of restriction enzymes and molecular biology reagents.

### Data availability

All raw and processed sequencing data generated in this study have been submitted to the NCBI Gene Expression Omnibus (GEO; https://www.ncbi.nlm.nih.gov/geo/) under accession number XX.

## Materials

All human FFPE genomic DNA used in this study was obtained from BioChain Institute, Inc. (Newark, CA). All enzymes used in this study were obtained from NEB Inc. The DNA repair step commonly used in this manuscript corresponds to the NEBNext FFPE DNA Repair v2 Module and contains enzymes and buffers that are optimized to repair FFPE DNA in next generation sequencing workflow including UDG, Endonuclease IV, Fpg, Endonuclease VIII, T4 PDG, Taq DNA ligase and Bst DNA polymerase.

### Uracil detection from FFPE human genomic DNAs without and with DNA repair

Three DNA polymerases, Q5 DNA polymerase (Q5), Q5 dU-bypass DNA polymerase (Q5U, for which a mutation in the uracil-binding pocket enables the enzyme ability to read and amplify templates containing uracil and inosine bases) and Taq DNA polymerase (Taq) were used to evaluate uracil distribution present in FFPE DNA sample. Each condition was prepared with and without DNA repair. Human FFPE genomic DNA from lung tumor tissue was fragmented to 200 bp average size using a Covaris S2 (Covaris Inc.) with the following settings: 10% duty cycle, intensity 5, 200 cycles per burst and treatment time of 6 minutes. 50 ng of fragmented DNA was used for each library preparation. Without DNA repair, libraries were constructed using NEBNext Ultra II DNA Library Prep Kit for Illumina (NEB, Inc.) with these modifications. First, Illumina Y adaptor was used to replace NEBNext loop-adaptor. Second, no USER treatment after adaptor ligation, and finally, PCR enrichment was performed using Q5 DNA polymerase, Q5 dU-bypass or Taq DNA polymerase respectively to amplify the adaptor-ligated libraries. With DNA repair, prior to the end repair step, the fragmented DNAs were treated with FFPE DNA repair module v2 at 20°C for 30 minutes. All other steps in the library preparation protocol were kept the same as for the unrepaired DNA library construction. Libraries were prepared in duplicate for each condition. The library quality was assessed using a high sensitivity DNA chip on a Bioanalyzer (Agilent Technologies, Inc.). All libraries were indexed and paired-end sequenced on an Illumina MiSeq platform.

#### Purification of 5mC deamination repair enzymes

Human thymine-DNA glycosylase (hTDG WT and hTDG-A145G) was cloned into pET28-based expression vector with 6-his tag at the C-terminus of the protein. MBD4 was cloned into pET28-based expression vector as a fusion of 14-His-MBP-SUMO at the N-terminus of MBD4. Both constructs were transformed into NEB T7 Express cells. Expression cells were grown in Luria broth at 37°C till A600nm = 0.6, following 0.4 mm isopropyl β-d-thiogalactoside induction at 18°C overnight. Cells were harvested and protein was purified as described below. For hTDG, crude supernatant (300 mM NaCl, 20 mM Tris, pH8, 1 mM DTT, 0.1 mM EDTA and 5% glycerol) was applied to a DEAE column. The Flow-through of DEAE column was diluted to final buffer containing 100 mM NaCl, 20 mM Tris, pH8, 0.33 mMDTT, 0.1 mM EDTA, 5% glycerol and was subjected to Heparin column, following Heparin HyperD (500 mM NaCl, 20 mM Tris, pH8, 0.1 mM EDTA, 5% glycerol), Ni-affinity (500 mM NaCL, 40 mM NaPO4, 250 mM Imidazole), and Source 15Q and dialyzed into storage buffer (200 mM NaCl, 20 mM Tris, pH 7.5, 1 mM DTT, 0.1 mM EDTA, 50% glycerol). For MBD4, crude lysate (300 mM NaCl, 20 mM Tris, pH8, 1 mM DTT, 0.1 mM EDTA and 5% glycerol) applied on DEAE, then HeparinHyperD, following with SenP1 cleavage 90 minutes at room temperature. The MBD4 cleavage product was further purified by a Source 15S column (elution at 168 mM NaCl, 20 mM Tris, pH 8, 1 mM DTT, 0.1 mM EDTA, 5% glycerol). Final purified protein was dialyzed into storage buffer (200 mM NaCl, 20 mM Tris, pH 7.5, 1 mM DTT, 0.1 mM EDTA, 50% glycerol). The purity of the proteins were greater than 99% assessed by SDS-PAGE with Coomassie staining. The concentration of the proteins was determined by absorbance at A280.

### NGS assay to determine the efficiency of enzymatic repair on 5-methylcytosine and 5-hydroxymethylcytosine deamination

Bacteriophage XP12 and T4gt genomic DNA was used to generate 5-methylcystosine and 5-hydroxymethylcytosine deamination substrates by heat alkaline treatment. 1500 ng of phage genomic DNA was diluted to 30 ng/ul using 1xTE buffer and sheared to 200bp using Covaris S2 standard protocol. 125 ng of fragmented DNA was used to prepare each DNA library following the NEBNext Ultra II library prep kit for Illumina (NEB, Inc.) until the loop-adaptor ligation step. After ligation, the sample was subjected to heat alkaline deamination by adding an equal amount of 1M NaOH into adaptor-ligated DNA following incubation at 60°C for 20 minutes. An equal amount of 1M acetic acid was added to neutralize the alkaline treated sample immediately. The neutralized DNA sample was further purified using Zymo oligo DNA clean and concentrator kit (Zymo Research). The Zymo-cleaned DNA was subjected to anneal by incubating at 90°C for 2 minutes following gradually cooling down to 20°C. This 20 minutes deaminated re-annealed XP12 or T4gt gDNA was subject to USER treatment only for non-repair sample; for repair sample, hTDG-A145G or MBD4 (0.25 uM) was also added with USER enzymes, incubated the reaction at 37°C for 60 minutes. All enzymes were removed by thermolabile proteinase K at 37°C for 15 minutes following 60°C for 15 minutes. PCR amplification was performed according to the manufacturer’s protocol. All the libraries were evaluated using a high sensitivity DNA chip on a Bioanalyzer (Agilent Technologies). All libraries were indexed and paired-end sequenced on an Illumina MiSeq platform.

### Deep sequencing of cancer gene panel from FFPE human lung DNA

FFPE human lung genomic DNAs were diluted in 0.1x TE buffer and sheared to 200bp using a Covaris S2 as described earlier. 200 ng fragmented FFPE human lung genomic DNA was subjected to library preparation following the NEBNext Ultra II DNA Library Prep workflow without and with DNA repair. Without DNA repair, 200 ng fragmented DNA was blunted and dA tailed followed by adaptor ligation. With DNA repair, prior to DNA blunting and dA addition step, 200 ng fragmented DNA was treated with 1) NEBNext FFPE DNA Repair v2 Module; 2) hTDG; 3) NEBNext FFPE DNA Repair v2 Module plus hTDG, incubated at 37°C for 30 minutes followed by end prep step. Pre-capture libraries were PCR amplified with NEBNext Multiplex PCR primers and Ultra II Q5 Master Mix for 8 cycles. Libraries were quantified on an Agilent Bioanalyzer, and 750 ng libraries were used for target capture with Agilent XT ClearSeq Comprehensive Cancer Panel. Post-capture libraries were amplified with Ultra II Q5 Master Mix for 14 cycles. Libraries were pooled and sequenced on an Illumina MiSeq at 2 × 75 bp. Data from a total of four Miseq runs were combined and analyzed for variant calling.

## Data analysis

### Read trimming and mapping

Paired-end Reads were trimmed using Trim Galore (version 0.6.7) with the following parameters (--clip_R1 1 --clip_R2 1 --three_prime_clip_R1 1 --three_prime_clip_R2 1 --paired). Trimmed reads were mapped to the human genome (hg19) using bwa mem (default parameters). For the XP12 and T4gt experiments, trimmed reads were mapped to the phage XP12 (MT664984.1) and phage T4 reference genome (NC_000866.4) respectively using bwa mem (default parameters).

### Overall fraction of CG:TA calculation

The mapped reads are processed using the damage_estimator program (from https://github.com/Ettwiller/Damage-estimator) (3) More specifically, mapped Read1 and Read2 from paired-end reads are separated into two files using the split_mapped_reads.pl using the default parameters. The resulting mpileup files are analyzed using estimate_damage_location.pl and visualized in R using the following programs : plot_damage_location.R or plot_damage_location_context.R. For the low, medium and high frequency variants, the Read1 were used only and the corresponding mpileup files obtained using split_mapped_reads.pl were downsampled to 300 fold coverage per-position. Positions with less than 300 were discarded. Positions that have low (<=1%, 3 reads or less) medium (1%-10% 4 to 30 reads) and high frequency variants (>10%, 31 reads or more) were counted and normalized to all the positions to obtain the number of low/medium and high frequency variants per megabase sequence. Only read variants with base quality phred score above 30 were used.

